# PALINCODE: Recording cell lineage with ternary palindromic CRISPR bits

**DOI:** 10.64898/2026.04.16.718941

**Authors:** Maryam Fathi, Aidan Cook, Bagheri Meisam, Tyler Curiel, Aaron McKenna

**Affiliations:** Department of Molecular and Systems Biology, Geisel School of Medicine, Dartmouth College, Hanover, NH; Dartmouth Cancer Center, Lebanon, NH; Graduate Program in Quantitative Biomedical Sciences, Geisel School of Medicine, Dartmouth College, Hanover, NH; Program in Pharmacology, Boston University, Boston, MA

## Abstract

Reconstructing complete and accurate lineage trees remains a long-standing challenge in biology. Here, we introduce PALINCODE (**Palin**dromic **Co**ding and **De**coding), a system that utilizes ternary CRISPR bits (cBits) to stochastically write one of three possible states over time, permanently embedding lineage relationships in the genome. We demonstrate PALINCODE’s lineage-recording potential through simulations and establish palindromic CRISPR editing in cell culture models. We show that truncated Cas9 guide sequences yield ternary outcomes at high efficiency when compared to conventional guides. Using PALINCODE, we derived lineage-recording cell lines with a theoretical coding capacity of up to 10^25 bits, enabling the generation of lineage trees 32 cell divisions deep in single-cell sequencing of 293T cells. Furthermore, we applied PALINCODE using an in vivo melanoma model to jointly read out lineage history and gene expression, enabling in vivo reconstruction of clonal evolution within tumor cell clonal populations. PALINCODE circumvents several limitations of prior CRISPR-based systems while increasing the information potential at individual CRISPR sites, creating a lineage-recording platform with higher density than many competing approaches.

## Introduction

CRISPR lineage tracing technologies have rapidly evolved in both complexity and biocompatibility. New genome-editing strategies, such as base and prime editing, have overcome many challenges posed by first-generation CRISPR systems that relied on cytotoxic double-stranded breaks (Choi et al. 2022; Winter et al. 2025; McKenna and Gagnon 2019). Prime-editing systems can generate diverse editing outcomes at each locus and have been utilized in both sequencing and spatial lineage systems (Koblan et al. 2025; Choi et al. 2022). Conversely, base-editing systems offer lower diversity in editing outcomes but benefit from straightforward engineering and compatibility with both sequencing and spatial profiling approaches (Winter et al. 2025; Chadly et al. 2024).

These technologies highlight an ongoing challenge: the need for efficient, high-resolution lineage-tracing systems that are biologically tractable but can generate stochastic editing outcomes. To address this, we developed PALINCODE (**Palin**dromic **Co**ding and **De**coding), a CRISPR-based lineage recording system. PALINCODE introduces a novel strategy for recording lineage using palindromic CRISPR target sites. PALINCODE target sites can be edited in one of two mutually exclusive orientations (left or right). This approach preserves the stochastic nature of earlier indel-based CRISPR systems while leveraging the simplicity of base-editing approaches.

To validate PALINCODE’s performance, we applied it across multiple experimental systems. In vitro, we introduced PALINCODE constructs into human 293T cells to assess editing behavior and lineage encoding under controlled conditions. For in vivo validation, we transplanted A375 cells containing integrated PALINCODE barcodes into immunocompromised mice to track tumor formation. By combining a scalable delivery strategy with biologically compatible, DSB-free editing, PALINCODE demonstrates robust performance in both cultured cells and complex in vivo models, offering a practical framework for building high-resolution lineage maps in experimental and therapeutic settings.

## Results

We sought to develop a compact lineage-tracing system using CRISPR/Cas9 base editing that preserves the stochastic nature of our previous recording systems while minimizing information loss due to double-stranded breaks(Raj et al. 2020; McKenna and Gagnon 2019; Simeonov et al. 2021; McKenna et al. 2016). We envisioned a system where states are randomly selected from a mutually exclusive set of outcomes and irreversibly recorded in cellular DNA. While overlapping guides can achieve this by jointly editing a single target site, balancing the efficiency of two guides is challenging and often requires expressing multiple guide sequences (Farzadfard et al. 2019).

Here we employed palindromic CRISPR target sites, allowing the Cas9 base-editing enzyme to bind to either strand of DNA. In this design, base editing at the distal end of the target occurs in the ‘seed’ region of the opposite orientation, preventing subsequent editing in the opposing direction (**Figure 1A**).

**Figure 1:**
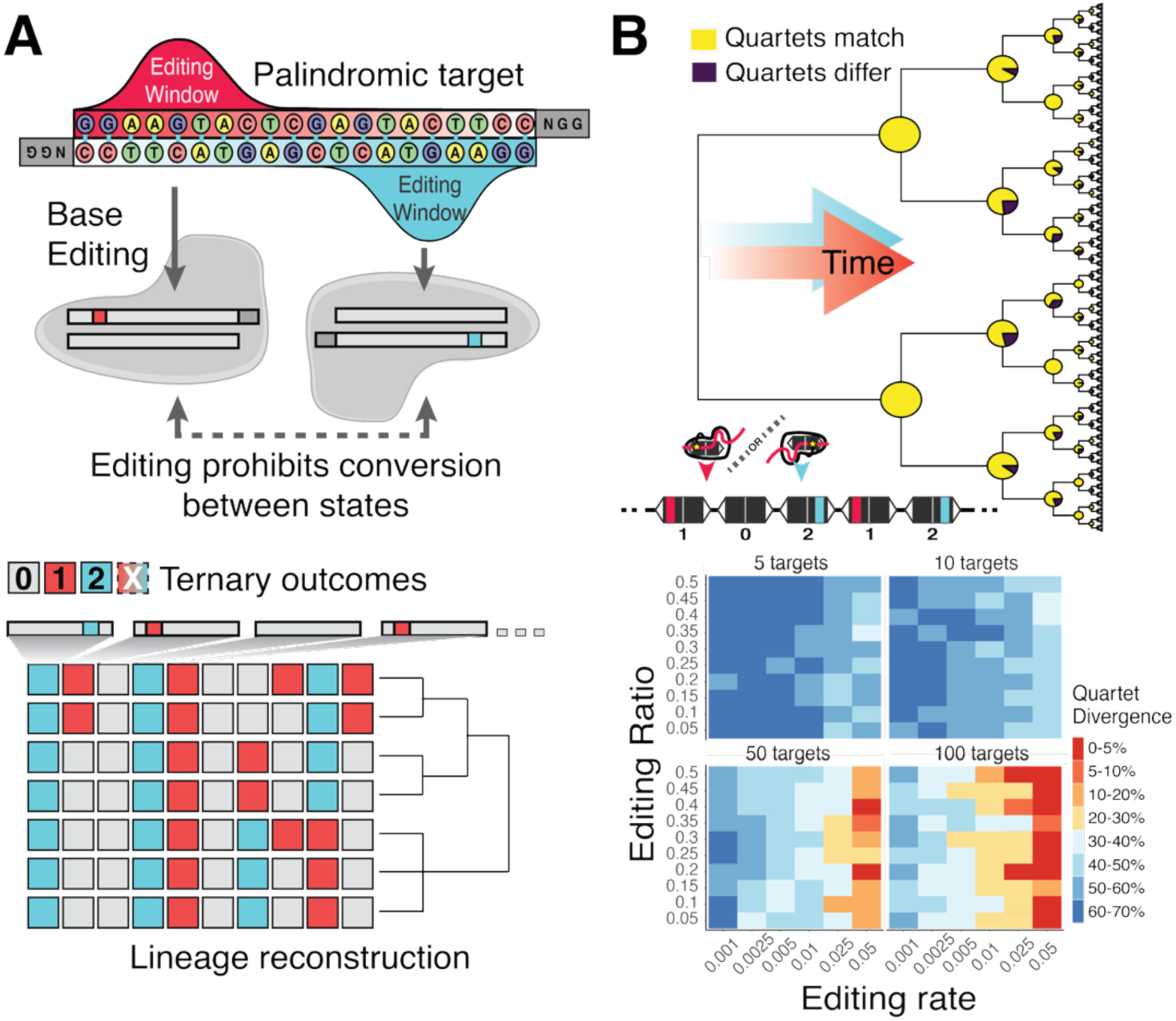
PALINCODE CRISPR ternary bits (cBits). **(A)** The PALINCODE system relies on palindromic CRISPR targets (cBits), where a core sequence is mirrored on the top and bottom strand with the protospacer adjacent motif (PAM) terminating each end. CRISPR-Cas9 deaminase can then bind in either orientation, converting a distal adenine within the editing window to guanine. This guanine sits in the core binding region of the opposite strand, preventing base editing in that orientation. These ternary outcomes can then be collected for each cell, and lineage histories can be reconstructed**. (B)** Simulations of a dividing population of cells encoded with ternary cBits, measured with quartet divergence. By varying the editing balance between left and right outcomes, the overall editing rate per cellular generation, and the number of integrated targets, we can identify parameters that minimize quartet divergence.

This approach creates a ternary cBit (trit): the ‘0’ state represents the wild-type sequence, while states ‘1’ and ‘2’ represent exclusive editing on the forward or reverse strand, respectively, all derived from a single CRISPR guide and target site. While Chow et al. established the foundation for trit encoding using recombinases, CRISPR sites offer a more flexible and significantly more compact recording approach (Chow et al. 2021). PALINCODE encodes ternary outcomes at a density approximately an order of magnitude higher, with tunable rates that are driven by the target sequence.

To assess the accuracy of lineage reconstruction with this system, we simulated reconstruction capacity across a wide range of parameters, varying (1) the editing rate, (2) the imbalance between ternary outcomes, and (3) the number of CRISPR target sites (**Figure 1B**). We evaluated accuracy using the quartet distance (Smith 2022), a measure of the correct arrangement of every possible set of four leaves on the lineage tree. Recorders with 30 target cBits and well-balanced outcomes produced highly accurate trees (5–10% divergence from ground truth, **Supplemental Figure 1**). Increasing to 50 recorders yielded <5% divergence across a range of left-right balances, provided sufficient Cas9 editing rates were achieved (**Figure 1C, Supplemental Figure 1**).

### Establishing Palindromic CRISPR Editing

Guided by our simulations, we optimized experimental conditions to enable *in vivo* recording with palindromic targets. We selected two conventional genomic CRISPR targets, T7 and RNF2, which were previously characterized for base-editing efficiency. We engineered palindromic versions by mirroring the first 10 bases in reverse complement for bases 10–20 (**Supplemental Table 1**)(Kim et al. 2019). These targets were then integrated into the genome using the PiggyBac system.

While non-palindromic targets readily accumulated base edits using the CRISPR adenine base-editing construct ABEMAX, the palindromic versions showed poor efficiency (**Figure 2A**)(Koblan et al. 2018). Non-palindromic targets displayed adenine-to-guanine conversions within a window centered five base pairs from the distal end relative to the PAM (**Figure 2B**). In contrast, palindromic targets showed limited editing, though edits were observed in windows on both sides of the target sequence (**Figure 2C**). Computational scoring of palindromic target efficiencies using standard on-target metrics did not indicate that the target sequences were poor candidates (McKenna and Shendure 2018).

**Figure 2:**
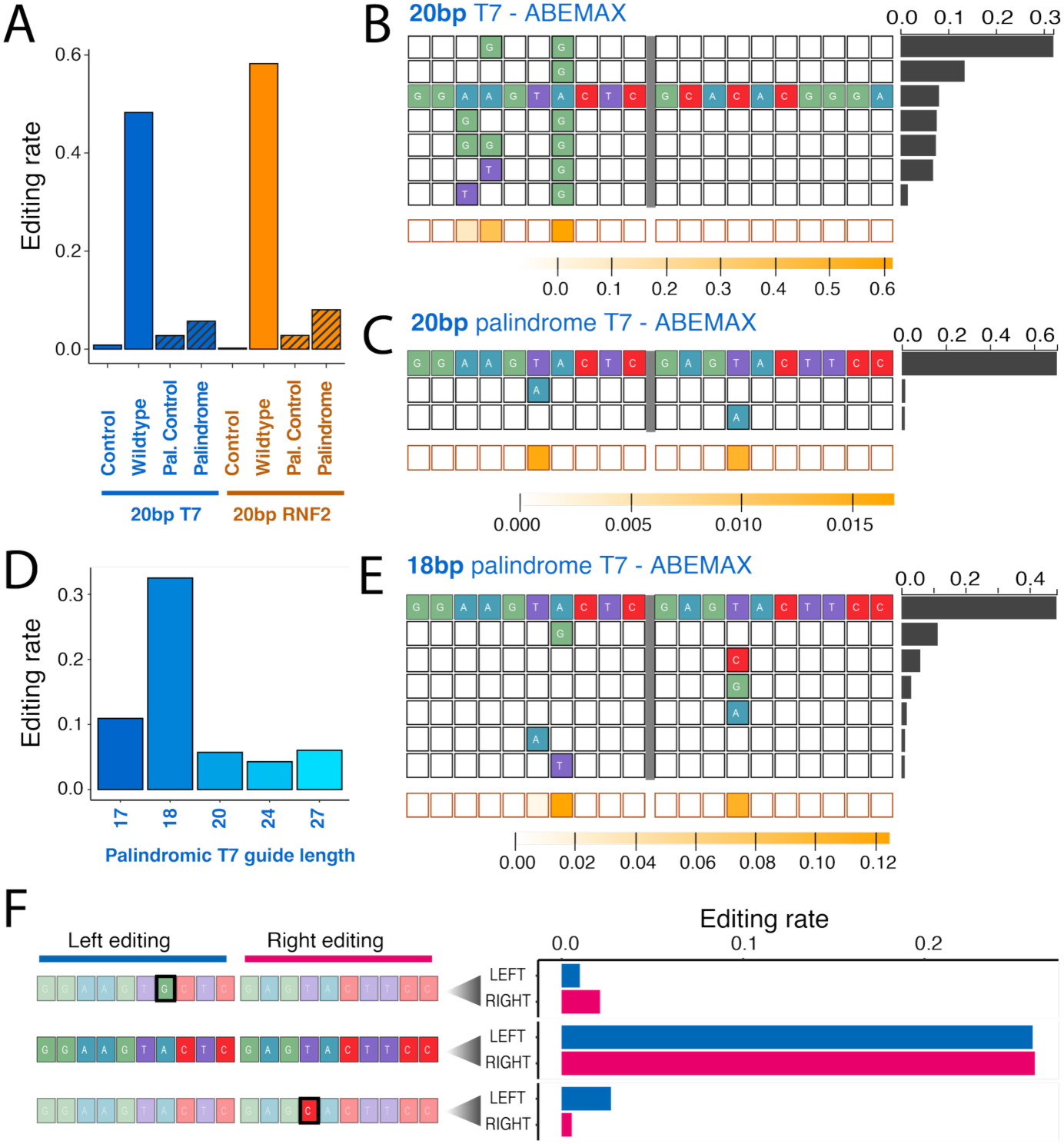
Guide sequence length determines editing efficiency and orientation. **(A)** Editing efficiencies using the 20 nucleotide T7 and RNF2 target and guide sequences compared to both palindromic versions, PalT7 and PalRNF2, as well as the control (no guide) in 293T cells. **(B)** Editing outcomes for a 20bp guide targeting the endogenous T7 target site in 293T cells, with editing occurring within the canonical window in various combinations at >90%. The histogram on the right indicates the proportion of each outcome, and the orange boxes at the bottom indicate overall editing per position. **(C)** 20 nucleotide palindromic T7 (PalT7) guide, where canonical and opposing editing occur at low frequency. **(D)** Editing across a range of palindromic PalT7 guide lengths, with editing increasing at the shorter guide length of 18. **(E)** An 18-basepair palindrome guide sequence targeting the 20-basepair target site, showing the increase in editing efficiency and mutually exclusive editing outcomes (left or right). **(F)** PalT7 target sequences were created with pre-existing left or right single-base edits, pooled with the wild-type sequence, and transfected with ABEMAX in a single cell-culture pool. Pre-edited sequences showed lower editing rates than the wild-type sequence.

We hypothesized that the secondary structure of palindromic guide sequences might inhibit CRISPR editing. We observed that folding energy scaled with guide length, as shortening the palindromic guides reduced the resulting hairpin size (**Supplemental Figure 2A, B, Supplemental Table 2**). We then investigated the use of guide sequences of different lengths to overcome editing issues, as previous studies have indicated that both longer and shorter Cas9 guide sequences can edit their genomic targets (Ran et al. 2013; Fu et al. 2014). We tested 17-, 18-, 20-, 24-, and 27-nucleotide guides against our PalT7 target sequence using the ABEMAX system (**Figure 2D**). Truncated guides (17 and 18 bp) significantly outperformed longer sequences, with the 18-nucleotide guide editing over 30% of target sequences. Similarly, for the palindromic PalRNF2 guide, the 18-nucleotide sequence yielded higher editing rates than the 20-nucleotide version, though overall efficiency was lower than that of the PalT7 target.

We next compared ABE8e to the ABEMAX editor. For the palindromic PalRNF2 target, ABE8e increased editing rates with the 20-nucleotide guide but resulted in a loss of the mutually exclusive editing pattern (**Supplemental Fig. 2C**). However, by switching to 18-basepair guides with ABE8e, we recovered left-right exclusivity (**Supplemental Fig. 2D**). We observed generally exclusive editing on both sides of the recorder; the probability of this dual-editing event was proportional to the joint probability of independent side editing, suggesting it arises from simultaneous binding events rather than sequential rounds of editing.

To confirm that left- or right-editing effectively inhibits editing in the opposite orientation, we tested targets with pre-engineered nucleotide changes corresponding to the most prevalent left- or right-edited states (**Figure 2F**). We observed an order of magnitude less editing compared to the wild-type sequence In the pre-edited palindromic PalT7 targets (average of 2.4% in pre-edited vs 26% in unedited) and in the PalRNF2 target (8% and vs 22%, **Supplemental Fig. 2E**). This suggests that target sequences remain locked in their state after the initial editing event and any editing on both sides of the target is most likely a simultaneous event.

### Determinants of palindromic editing outcomes

We next wanted to determine whether the observations in our PalT7 and PalRNF2 guides and targets would be consistent across a larger set of sequences, and which sequence features were driving palindromic target editing. We designed 208 palindromic target sequences with balanced GC content and varying numbers of adenines within the known editing window (**Figure 3A**, **Supplemental Table 3**). We synthesized and integrated this library into 293T cells via the PiggyBac transposase system. Flow-sorted cells were transfected with ABEMAX or ABE8e, and DNA recorders were captured 14 days post-transfection.

**Figure 3:**
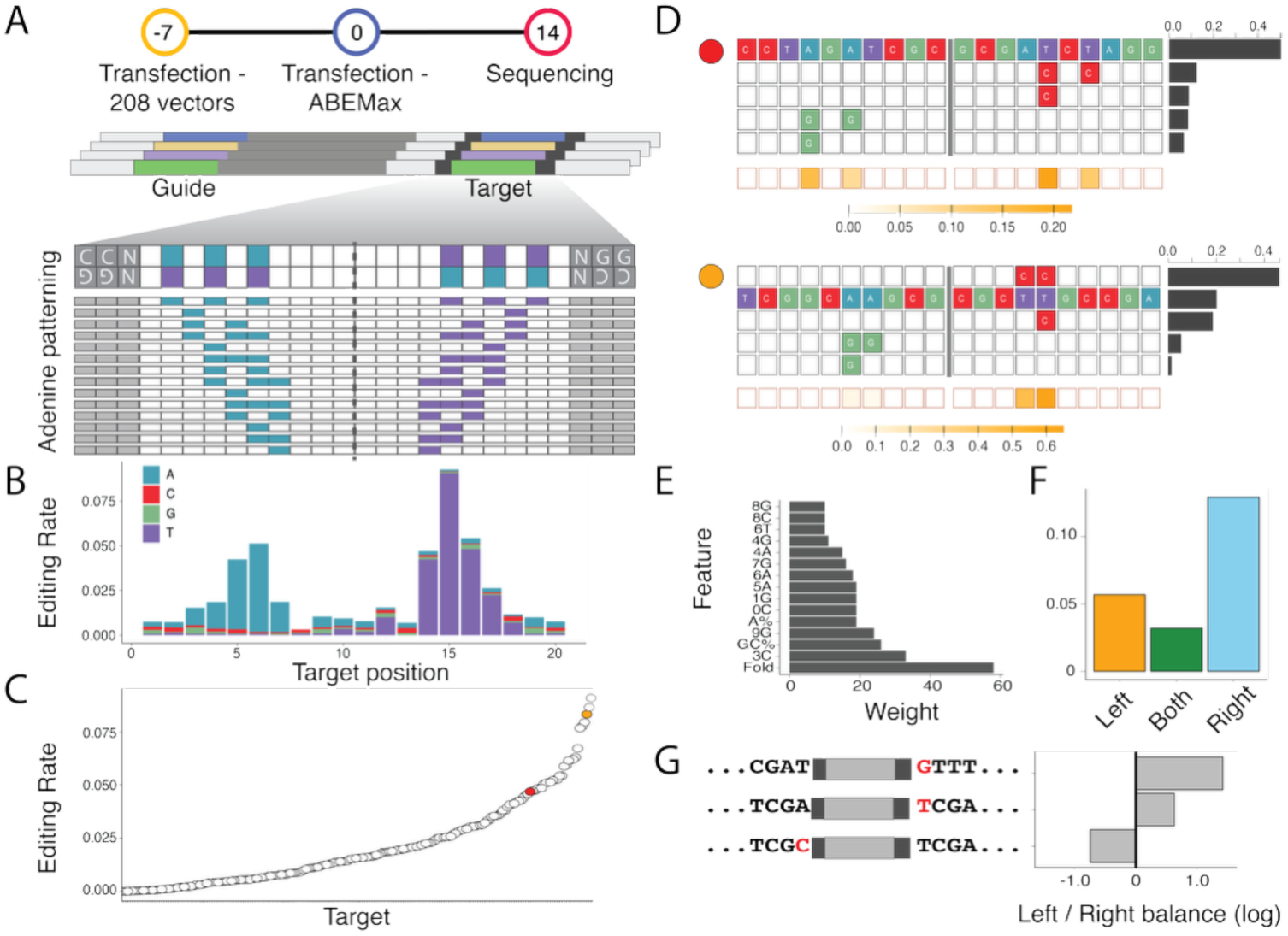
Determining key modulators of PALINCODE activity. **(A)** 208 PALINCODE guide-target pairs were created with varying numbers and patterns of adenines within the editing window in both orientations (bottom). These targets and guides were engineered into 293T cells using the PiggyBac transposase system. After 7 days of selection, Cas9 base editors were transfected and selected for an additional 7 days, after which targets were sequenced. **(B)** Aggregated base-editing outcomes across all targets, in either left- or right-handed orientations, show enrichment for right-hand editing. **(C)** Editing rates for individual targets varied widely over the 208 target sequences. **(D)** Two highlighted outcomes, matching colored dots in panel C, show targets with a relatively well-balanced distribution between left and right outcomes (top) and a more unbalanced target (bottom). **(E)** Extracted features ranked by weight show that folding energy is a critical determinant of efficiency in palindromic targets (Fold = folding energy, GC = GC percentage of target, 8C=cytosine at position 8, etc). **(F)** Left-right editing outcome balance in the original pool of CRISPR targets and **(G)** altered proximal (+1 bases) and their effect on left-right editing balance.

Editing was abundant, with enrichment at programmed adenines within the target sequence when using 18-nucleotide guides (**Figure 3B**). Compared to 17- and 20-nucleotide guides, the 18-nucleotide guides increased editing at adenines while minimizing dual-sided editing, particularly with ABE8e (**Supplemental Figure 3A, B**). Editing rates correlated well between ABEMAX and ABE8e editors (**Supplemental Figure 3C**), though individual targets exhibited a wide efficiency range (∼0% to ∼60%) (**Figure 3C**). Within the oligo target pool, the balance between left and right editing rates varied (**Figure 3D, E**), with some targets presenting balanced left and right outcomes, though many favored the right-hand orientation for editing.

We next modeled the key factors driving efficient editing across the target set. While conventional CRISPR scoring metrics (e.g., nucleotide content, GC proportion) were included(Konstantakos et al. 2022), the folding energy of the guide emerged as the dominant predictor of editing efficiency for palindromic sequences (**Figure 3E**), mirroring our results with PalT7 and PalRNF2 targets.

We were also curious what was driving the observed imbalance favoring editing on the right side of the target sequence (**Figure 3E**). Given that the sequence is identical in either orientation (including PAMs), we hypothesized that proximal bases outside the target influenced editing orientation.

Previous prediction tools highlight the importance of a guanine at the +1 position(Doench et al. 2014). To test this, we created a parallel CRISPR palindrome target library varying the first base outside the PAM on both ends (G on right only, neither, or left only) (**Figure 3G**). Editing balance shifted to match the orientation of the G nucleotide as seen by the bound CRISPR enzyme, providing a method to tune the balance of future palindromic CRISPR targets.

### Single-cell lineage reconstruction in cell culture

To validate PALINCODE as a tool for single-cell information encoding, we integrated a recorder construct containing the PalT7 and PalRNF2 targets, each driven by the U6 polymerase III promoter (**Figure 4A**). Each recorder construct contained a static “integration barcode”, a unique 12-nucleotide identifier marking the genomic integration site. Guides were also expressed via the U6 promoter, and the complete construct was integrated into 293T cells through successive rounds of transfection and flow sorting. We then isolated single-cell clones and selected one (Clone 9) for deep characterization. DNA sequencing of Clone 9 revealed 79 stably integrated recorders, representing 158 potentially editable target sequences. Progeny of this clone was transduced with CRISPR-CBE8e, single-cell sorted into individual wells, and expanded for both lineage characterization and single-cell sequencing (**Figure 4A**).

**Figure 4:**
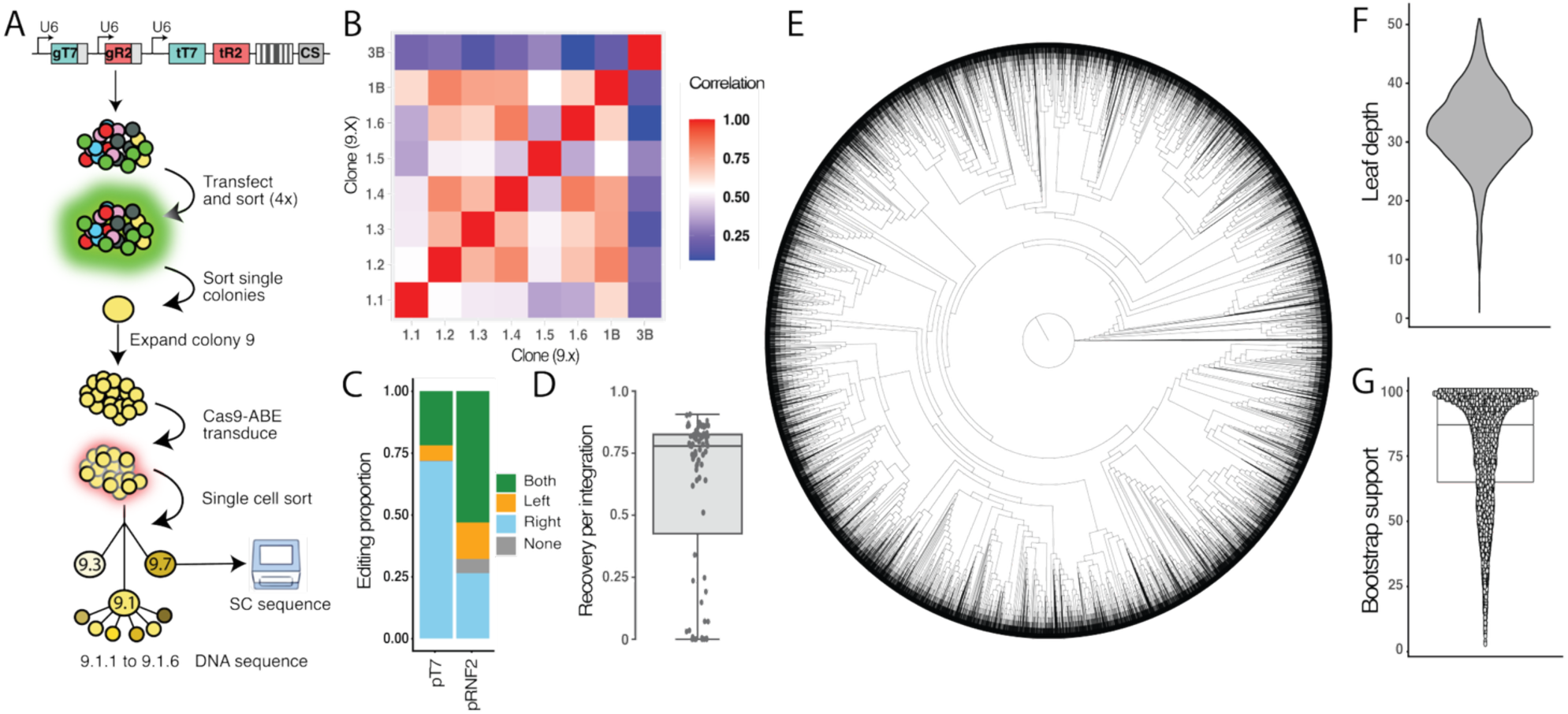
Single-cell lineage tracing with PALINCODE in 293T cells. **(A)** PalT7 and PalRNF2 guide sequences are individually expressed under U6 promoters, coupled with expression of their cognate targets and a static identifier under the expression of a U6 promoter. 293T cells were then transfected, sorted, and single-cell expanded into clonal populations. **(B)** The correlation of DNA barcode readout between subpopulations in clonal population 9 (3B, bulk population 9.3, 1B= bulk 9.1, 1.1 to 1.6=single-cell isolated populations from clone 9). **(C)** Editing outcome for palindromic PalT7 and PalRNF2 targets across single-cell sequencing of clone 9.7 and **(D)** average recovery of the 79 lineage recorders across the 7033 cells captured. **(E)** The resulting lineage tree was assembled with IQTree2 with an average leaf depth of 32 and **(F)** average bootstrapping support of 87% **(G)**.

We first analyzed the relationship between the Clone 9.1 parent population, its bulk subclonal populations (9.1.1 to 9.1.6), and an independent outgroup (9.3) (**Figure 4A, bottom**). Average editing outcomes for the 158 target sequences (78 x 2 per integration) were calculated from bulk DNA. The resulting correlation structure (**Figure 4B**) recapitulated known clonal relationships: 9.1 clones shared a strong lineage signal with the parent population, whereas the 9.3 outgroup showed little to no correlation.

To achieve higher resolution, we performed single-cell sequencing on an isolated subclone (Clone 9.7). Cells were processed for both lineage barcodes and transcriptional profiles using the 10x Chromium platform and lineage barcode enrichment (**methods**), yielding 7,033 single-cell profiles. Lineage recording activity was high, with fewer than 5% of sites unedited across all integrations (**Figure 4C**). When aligning lineage data with single-cell profiles, we observed variable recovery for the 79 DNA-validated integrations, with a median integration capture rate of 78% (**Figure 4D**). The loss or low expression of some recorders suggests transcriptional suppression or silencing for some integrations.

We then utilized the lineage recordings to construct a single-cell lineage tree. PALINCODE ternary outcomes were one-hot encoded into a binary matrix, and IQTree2 was used to reconstruct the phylogeny (**Figure 4E**)(Minh et al. 2020). The resulting tree showed a median cell-division depth of 32 and a maximum depth of 51 (**Figure 4F**). Bootstrapping indicated strong support for most branches, with a median support value of 87% (**Figure 4G**). This demonstrates PALINCODE’s ability to generate deep single-cell lineage trees using off-the-shelf single-cell sequencing technologies.

### Profiling evolutionary selection in melanoma

To study tumor evolution in vivo with high temporal and clonal resolution, we employed a xenograft transplant model using the human melanoma cell line A375. A375 cells harbor a BRAF V600E mutation and reproducibly form tumors in immunocompromised mice, enabling controlled interrogation of evolutionary dynamics. We engineered A375 cells using successive rounds of integration (methods) with the PALINCODE system, followed by a final transfection with the ABE8e editing construct. Each cell represents a single clone, tagged by a number of integration IDs carried by the PALINCODE lineage recorder (**Figure 5A**). The resulting cells were transplanted them into mice, and following tumor expansion, we utilized single-cell sequencing to jointly recover lineage histories and transcriptional states.

**Figure 5:**
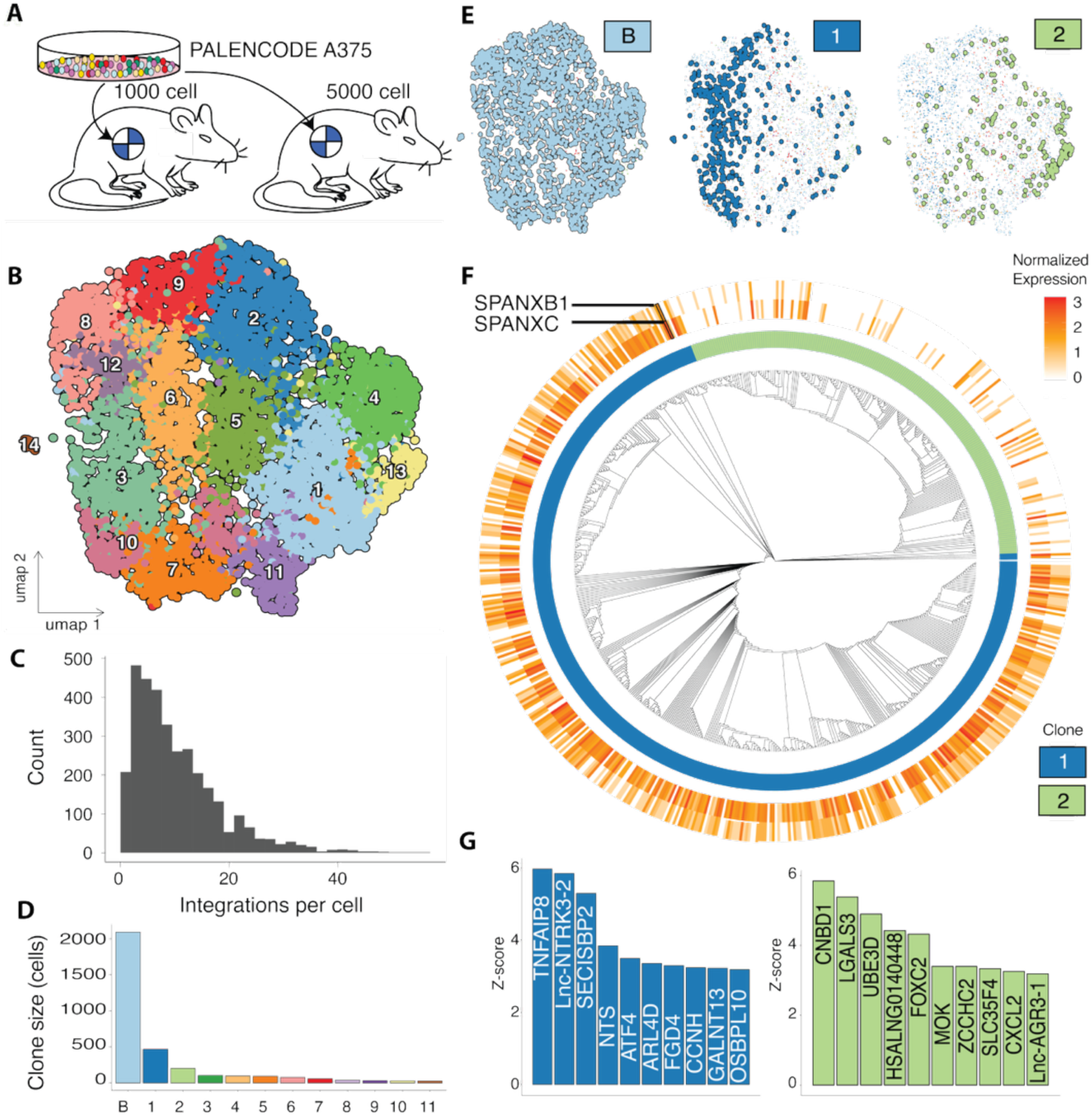
Single-cell lineage tracing in an A375 melanoma model. **(A)** A population of A375 cells was engineered with successive rounds of PALINCODE barcodes, creating a diverse population of clonal cells, which were activated with CRISPR-ABE and injected into two replicate mice. **(B)** After contaminating mouse cells were removed, the remaining 5,715 cells were arranged into 14 clusters. **(C)** When integrated with lineage recorders, 3,327 cells had at least one recorder, with a median of 8 integrations per cell in lineage+ cells. **(D)** 1,236 A375 were clustered into 11 clonal populations, with clonal sizes ranging from 1 to 1,236 cells (B = background clones, clones 1, 2, etc). **(E)** Clones 1 and 2 compared to the background clones (B) within UMAP space. **(F)** Recovered tree structure for clones 1 and 2, with the expression of SPANXB1 (outer) and SPANXC (inner ring) in individual cells. **(G)** PATH Z-scores of the top 10 most heritable genes on the resulting lineage trees for clones 1 and 2.

We recovered cells from four tumor sections across two mice, all initiated from a common population of PALINCODE A375 cells (**Figure 5A**). After 10X Chromium sequence and removal of contaminating mouse cells, the remaining 5,715 human cells were clustered into 14 populations (**Figure 5B**). These clusters were defined by the expression of known melanoma marker genes, including *SOX5*, *CPA4*, *S100A6*, and *LRP1B* (**Supplemental Table 4**). We then matched

PALINCODE lineage sequencing to cell identifiers captured in the single-cell sequencing. Of the 5,715 single cells, 3,327 (58%) contained at least one integrated lineage recording. In these cells, we captured a median of 8 integrated lineage recorders (**Figure 5C, Supplemental Figure 5B**). To define clonal populations within the diverse A375 starting pool, we clustered cells sharing static integration barcodes (methods). Of the 3,327 lineage-containing cells, 2,091 (62%) belonged to a diverse background of small clones with insufficient overlap to cluster cohesively. The remaining 1,236 cells (38%) clustered into 11 distinct clones, dominated by several large, expanding populations (**Figure 5D**). The two largest populations (Clone 1 and Clone 2), comprising 672 cells, were separated in the low-dimensional embedding within the background of other A375 cells (**Figure 5E**).

To dissect the mechanisms of tumor evolution in these competing clonal populations, we first performed differential expression analysis between each clone and the background of the smaller clones (**Supplemental table 5**). Both clones expressed a differing list of melanoma signature genes, including interferon and immune response genes (clone 1, *ISG15, IFI6, IFI27, LY6E, C3*), invasion (Clone 2, *CAV1, CDH13, CCN1*), and tumor suppressors and growth factors (Clone 1, loss of *LRP1B*; clone 2, upregulation of *IGFBP7*). Clone 1 was also enriched for Ki-67 expression when compared to other clones (**Supplemental table 5)**. Comparing Clones 1 and 2 revealed high expression of *SPANXB1* and *SPANXC* in Clone 1 (**Figure 5F**), both known markers implicated in melanoma progression (Westbrook et al. 2004; Salemi et al. 2009).

We then turned to the structure of the resulting lineage trees for each clone. PATH heritability analysis revealed genes that are correlated with tree structure (**Figure 5G**)(Schiffman et al. 2024). In clone 1, this included *TNFAIP8*, a key modulator of stem cell state, and *ATF4*, a regulator of plasticity in melanoma (Tian et al. 2024; Meinert et al. 2024). Conversely, clone 2 showed high heritability for expression of *LGALS3* and *FOXC2*; *LGALS3* promotes melanoma progression by enhancing cell migration, invasion, and metastasis (Braeuer et al. 2012) and *FOXC2* drives epithelial-mesenchymal transition (EMT) in melanoma cells, leading to cancer stem cell-like properties in other cancer types (Mani et al. 2007). This highlights the ability of single-cell lineage tracing to capture cellular evolution both between and within cancer populations, and future efforts could explore the effects of treatment on this cellular state space.

## Discussion

Reconstructing complex multicellular histories requires recording technologies that are both information-dense and biologically unobtrusive. With PALINCODE, we present a lineage recording platform that balances these requirements by repurposing standard base editors to encode ternary information into palindromic CRISPR targets. A key advantage of this approach is its simplified engineering. Unlike prime editing systems, which require pegRNA design and often suffer from variable efficiency, or first-generation CRISPR double-stranded break events, PALINCODE utilizes established, highly active adenine base editors (Koblan et al. 2025; Choi et al. 2022; McKenna et al. 2016). By pairing these robust enzymes with standard, truncated guide RNAs, we achieved high-efficiency recording, significantly lowering the barrier to adoption for diverse experimental models.

A defining feature of PALINCODE is its ability to generate stochastic, mutually exclusive outcomes from a single cBit target site. While conventional base editing is often deterministic, converting a specific base to another with high predictability, our palindromic design enforces a ternary choice (wild-type, left-edited, or right-edited). This stochasticity is essential for resolving branching lineage trees, as it prevents the rapid saturation of signals observed in completely deterministic systems. Furthermore, this design is compact. Because a single genomic site functions as a ternary cBit (trit), PALINCODE achieves a higher recording density than recombinase-based approaches or sparse CRISPR barcode arrays (Chow et al. 2021). This compactness not only conserves genomic space but also renders the system theoretically compatible with spatial sequencing by hybridization.

The information density of PALINCODE translated into robust performance in cell culture and in vivo models. In 293T cells, we achieved high-resolution lineage trees with an average depth of 3 cell divisions, validating the system’s capacity to track extended proliferative histories. In our mouse melanoma model, the system successfully recovered clonal dynamics in vivo, distinguishing between metastatic-like clones expressing *SPANXB1* and invasive clones driven by *LGALS3* and *FOXC2*. This demonstrates that PALINCODE can effectively link lineage history to transcriptomic state to uncover the drivers of tumor heterogeneity, improving on existing lineage-tracing technologies for cancer (Simeonov et al. 2021; Saxe et al. 2025; Quinn et al. 2021).

Despite these capabilities, the use of palindromic sequences introduces specific technical challenges. Palindromic DNA has a strong propensity to form secondary structures, such as hairpins, which can complicate the synthesis and cloning of large-scale target arrays. Consequently, scaling the number of recorders per construct requires careful sequence design and assembly strategies.

Additionally, these secondary structures can induce polymerase slippage during PCR amplification and sequencing, leading to an elevated background error rate compared to non-palindromic targets, which we observe. While we mitigated this through computational filtering and unique molecular identifiers (UMIs), future iterations of the system will need to optimize library preparation protocols to minimize such artifacts.

Finally, the resolution of any stochastic recorder depends on the balance of its possible states. While we found that PAM-flanking guanine nucleotides can influence the left-versus-right editing bias, some targets exhibited unequal editing probabilities, which could reduce the maximum information entropy. Future refinements of PALINCODE will focus on precise tuning of the left-right balance through further guide-length optimization and flanking sequence engineering. The dual-editing outcomes (left and right) can also result from successive rounds of binding and editing or from a single event; our data suggest it is often a single event, but future efforts would need to explore this further. We have additionally built our trees using a simple outcome encoding that indicates which side of the recorder is edited, but in the future, more information could be recovered from individual nucleotide outcomes in the data, leading to deeper, more accurate lineage reconstructions. Lastly, our approach could be combined with other CRISPR editing approaches, including prime editing, to increase information content. By maximizing the entropy of each cBit (trit) and expanding the library of validated palindromic targets, PALINCODE offers a scalable, high-resolution framework for mapping the developmental and evolutionary histories of complex biological systems.

## Methods and Materials

### Plasmids Design

All constructs sequence maps for constructs are available on Addgene (**Supplemental Table 6**). All primers and gBlock DNA fragments were obtained from Integrated DNA Technologies (IDT). For plasmid construction, restriction enzymes, NEBuilder HiFi DNA Assembly Master Mix (#E2621) for Gibson assembly, and the Quick Ligation Kit (#M2200) were used, all sourced from New England Biolabs (NEB). Q5® High-Fidelity 2X Master Mix (#M0492) from NEB was used for PCR amplification. All PCR products and DNA fragments were purified using the Zymo Clean and Concentrator-5 Kit (#D4003). Plasmid vectors were purified using either the Qiagen Miniprep Kit (#27104) or the ZymoPURE II Midiprep Kit (#D4200).

### Base Editing Construction

#### pCMV-ABEmax-Puro-U6, ABE8e-Puro-U6

To be able to select for the transient expression of our base editors, we amplified the EF1 promoter and puromycin resistance gene from a PiggyBac plasmid and inserted them into the pCMV-ABE7.10, pCMV-ABEmax, ABE8e. In addition, we inserted the U6 promoter for gRNA expression with two distinct types of tracer RNA: Hsu et al. and Chen et al. tracer RNA for each base editor (Cameron et al. 2017; Chen et al. 2013).

#### pRDA-426-DsRed: A Constitutive dCas9-DsRed Expression Plasmid

We modified pRDA-426 (Addgene #179097) to make a fluorescent version (pRDA-426-DsRed) for stable, constitutive expression of d-Cas9 with DsRed. We amplified DsRed from EF.CMV.RFP vector (Addgene #17619) and cloned into the NotI-rSAP-pRDA-426 digested plasmid using Gibson assembly.

#### Twist Pool Construction

We modified the PiggyBac backbone and created a PiggyBac-BSD that is selected by Blasticidin. The U6 promoter was also cloned into PiggyBac-BSD to express the gRNA of our Twist array. We then designed the Twist array with a specific pair of primers at both ends, gRNA- scaffold- terminator-spacer-Palindrome target with dual PAM. The pool included different arrays of 17bp, 18bp, and 20bp gRNA. We amplified each array group using a specific pair of primers and NEB Q5 polymerase. Each group of the twist pool was digested with Esp3I and inserted separately into the Esp3I-digested Piggybac-BSD-U6 vector. We ligated each construct and then transformed it into NEB Stable Competent E. coli (#C3040). It was grown in 1ml of SOC and incubated for 1 hour at 37 °C on a shaker. Since it was a DNA library pool, we did not plate it, and after 1 hour incubation at 37 °C, 6ml LB, as well as 100 μg/mL Ampicillin, were added to the bacteria and incubated at 37°C overnight. The plasmid was extracted by Qiagen QIAprep Spin Miniprep Kit (#27106X4). We created the alternative PAM twist array by modifying our original twist array and changing the target overhang sequences and the PAM site. The cloning process was carried out using the same method as we described in the previous twist pool.

#### Construction of PiggyBac-BSD-PalT7-PalRNF2 gRNA Vector for Stable gRNA and UMI Expression

We derived the PiggyBac-BSD-PalT7-PalRNF2 gRNA from the blasticidin-resistant PiggyBac-BSD backbone. We designed it for stable, constitutive expression of gRNA with a UMI containing target, driven by distinct U6 promoters. We ordered PalT7 and PalRNF2 gRNAs with distinct U6 promoters as IDT gBlock and inserted them into SpeI-rSAP-digested PiggyBac-BSD plasmid by Gibson assembly. We transformed and extracted the construct plasmid as previously described.

#### Engineering a GFP-Labeled Palindromic Barcode Plasmid

To introduce GFP to our system, we ordered GFP as an iDT gBlock and used Gibson assembly to insert it into the SpeI-rSAP- PiggyBac-BSD-PalT7-PalRNF2 gRNA digested vector. Then, to clone the Unique Molecular Identifier (UMI) target, we ordered a single-stranded fragment (IDT 4 nmole Ultramer DNA oligo) and made it double-stranded by PCR. The double-stranded DNA fragment as well as PiggyBac-GFP-PalT7-PalRNF2 gRNA were digested by ESP3I. We used Quick Ligase for ligation. We transformed the resulting construct by electroporation into NEB^®^ 10-beta Electrocompetent *E. coli* #C3020K. We did electroporation using Bio-Rad Gene Pulser in a 1-mm gap cuvette (Cell Projects, #EP-101) at 1.8 kV, 200 Ω, and 25 μF. We transferred the cells into 500 µl of SOC medium and recovered for 1 h at 220 r.p.m. and 37 °C, and then added them to 30 ml Super Broth with Ampicillin at 100 *μ*g/mL for overnight growth. We purified the plasmid by Zymo midiprep #D4200 for high-diversity UMI plasmid production.

#### Pre-edited PalRNF2 and PalT7

We designed and ordered the pre-edited PalRNF2 and PalT7 targets as oligos with their own unique flanking identifier to distinguish each pre-condition (Left-edited, right-edited and wild-type “unedited”). We then annealed and phosphorylated each pair using NEB 10x T4 ligation buffer (#B0202) and NEB T4 PNK (#M0201), incubated 30 min at 37 °C, 5 min at 95 °C, and then ramped down to 25 °C at 5 °C/min. Each insert was separately dropped into the NcoI-EcoRI-rSAPed Piggybac-BSD and ligated. To ensure that the transfection or sequencing preparation did not bias our results, all three combinations were co-transfected with the base-editor into a single population of 293T cells and the resulting sequences were divided post-sequencing into separate bins for analysis based on the unique flanking tag.

#### Cell culture

We performed all tissue culture experiments with HEK293T (ATCC #CRL-3216) and A375 cells (ATCC #CRL-1619). We maintained HEK293T and A375 in complete DMEM, Dulbecco’s modified eagle medium (DMEM, Corning #10013CM) supplemented with 10% fetal bovine serum (FBS) (HyClone #SH30910.03), 100 units/mL penicillin and streptomycin (Corning #30002CI). Cells were cultured at 37°C in a humidified atmosphere containing 5% CO2. We maintained and split the cells by trypsinization HEK293T with 0.05% trypsin (Corning #25052CI), A375 with 0.25% (Corning #25053CI) of cells were passaged every 3-4 days as needed to avoid confluency.

#### Lentivirus Preparation

We seeded HEK293T in a 10 cm dish at a density of 4 × 10^6 cells in 10 mL of complete DMEM, 24 h before transfection. Transfection was performed using Lipofectamine™ 3000 transfection reagent (Invitrogen #L3000015). We combined 1500ul Opti-MEM (Gibco #31-985-070), 35ul P3000 reagent, 750 ng packaging plasmid pCMV_VSVG (Addgene #8454), 2500 ng psPAX2 (Addgene #12260), and 5000 ng of the expression plasmid (pRDA-8e-DsRed). We mixed this with 1500ul Opti-MEM and 41ul Lipofectamine 3000 and incubated for exactly 5 min at room temperature. We concentrated the lentiviral supernatant using Lenti-X™ Concentrator (Takara #631232) at a ratio of 1:3 ml. The supernatant was mixed gently and incubated at 4°C for 3 hours. We then centrifuged it at 1500g for 45 min at 4°C. The supernatant was removed and the pellet was resuspended in 500ul of Opti-MEM. The resuspended virus was then aliquoted into three tubes and stored at -80°C.

#### HEK293T Cell Line Engineering and Preparation for In Vivo Lineage Tracing

We engineered HEK293T with our cBit construct through co-transfection of transpose and 1200 ng of the barcode construct for constitutive expression using Lipofectamine™ 3000 transfection reagent according to the manufacturer’s protocol. Using fluorescence-activated cell sorting (FACS) on the Aria II, we collected the highest-GFP-fluorescence cells 24 hours after transfection. We subjected the sorted cells to two additional rounds of integration, resulting in a total of three rounds of transfection. We sorted cells with the highest GFP fluorescence after each transfection. Next, the sorted-high fluorescence barcoded cells were seeded in a 6-well plate with 3 ml of complete media at a density of 0.3 x 10^6 and incubated in a 37 °C incubator. The next day, we added 10μg/ml of polybrene and incubated for 2 hours at 37 °C. We infected our barcoded HEK293T cells by adding a dropwise tube of virus (the Cas9) to the cells. 72 hours after, we sorted the dual positive cells (GFP+ and DsRed+) and single-cell sort them in a 96-well plate.

#### Engineering HEK293T Cells with Pre-edited palRNF2 Constructs

We co-transfected HEK293T with 1200 ng of transposes and a mixture of the three pre-edited palRNF2 plasmid constructs (none, left, right edited each at 700 ng) using the Lipofectamine™ 3000 transfection reagent. We initiated the selection of the integrated cells 24 hours after transfection by adding 10 ug/ml blasticidin to the medium and maintained the cells under selection for 4 days before DNA collection, amplification, and sequencing.

#### Next-Generation Sequencing Library Preparations

We amplified the region of interest (the barcode) using primers listed in the supplementary material. Next, we amplified the PCR product with another primer pair to add an adaptor for the next-index PCR. For the final PCR and multiplexing, we used primers to specify each sample. We cleaned up each PCR reaction using AMPure beads.

#### Single-Cell RNA Sequencing of 293T Cells

293T were integrated with the cBit construct, and editing was activated with the ABE8e construct as detailed above. Cells were sorted on an Aria II to isolate single clones into individual wells of a 96-well plate. The resulting sequencing reads were processed with CellRanger 7.1.0. The filtered cellular IDs and their transcriptional data were then processed using the MULTI-seq pipeline in R, and further plotting and single-cell analysis were done in Scanpy or Seurat (McGinnis et al. 2019; Wolf et al. 2018; Satija et al. 2015).

#### A375 Cell Line Engineering and Preparation for In Vivo Lineage Tracing

We engineered A375 with our cBit construct through transfection of transpose and 1200 ng of the barcode construct for constitutive expression using Lipofectamine™ 3000 transfection reagent according to the manufacturer’s protocol. Using fluorescence-activated cell sorting (FACS) on the Aria II, we collected cells with the highest GFP fluorescence 24 hours after transfection. We subjected the sorted cells with three additional rounds of integration, resulting in a total of four rounds of transfection. We sorted cells with the highest GFP fluorescence after each transfection. Next, the sorted-high fluorescence barcoded cells were seeded in a 6-well plate with 3 ml of complete media at a density of 0.3 × 10^6 and incubated at 37 °C. The next day, we added 10μg/ml of polybrene and incubated for 2 hours at 37 °C. We infected our barcoded A375 cells by adding a tube of virus (the Cas9) to the cells. 72 hours later, we sorted the dual-positive cells (GFP+ and DsRed+) using FACS to enrich the barcoded cells. We also performed single-cell sorting into a 96-well plate to establish individual clones. We expanded these clones to analyze barcode editing efficiency, assess UMI overlap, and quantify barcode diversity. In parallel, we maintained the original bulk-sorted population for subcutaneous injections into immunocompromised mice to assess in vivo tumor formation and lineage tracing.

#### Mice

We used immunocompromised NSG® (005557) mice obtained from Jackson Laboratory. We housed all mice in pathogen-free conditions at the Center for Comparative Medicine and Research, Dartmouth Hitchcock Medical Center. We conducted all animal procedures in accordance with protocols approved by the Institutional Animal Care and Use Committee (IACUC) at Dartmouth College.

#### In Vivo Tumor Initiation via Subcutaneous Injection of A375 Engineered Cells

After four rounds of barcode transfection, we sorted for the cell population with the highest fluorescence using flow cytometry. We then infected these enriched cells with an ABE8e-expressing viral vector to enable constitutive dCas9 expression. Following two days of recovery and expansion, we subcutaneously injected the engineered cells into immunocompromised mice at varying doses (1,000 and 5,000 cells) to initiate tumor formation. To prepare the cells for injection, we resuspended each cell dose in a total volume of 50 μL, consisting of 10 μL Matrigel (Corning, Cat # 354234), the appropriate volume of cell suspension, and Opti-MEM to bring the volume to 50 μL. We monitored tumor growth every three days using caliper measurements. On day 20, we euthanized the mice in accordance with IACUC guidelines due to tumor size reaching the ethical endpoint. After dissection, we sectioned each tumor into four pieces and used pieces 1 and 4 for single-cell RNA sequencing analysis.

#### Single-Cell Transcriptomic Profiling of Barcoded Tumor Samples and In vitro HEK293T Cells

We performed single-cell RNA sequencing on both tumor-derived cells (from pieces 1 and 4 of the xenografts) and in vitro cultured engineered HEK293T cells. To process tumor samples, we enzymatically dissociated the tissue into single-cell suspensions using a digestion mix containing 1 mg/mL collagenase type I, 0.1 mg/mL DNase I, and 1 mg/mL dispase in RPMI-1640 medium. We incubated the mixture at 37 °C for 30–45 minutes with gentle agitation. After digestion, we filtered the suspension through a 40 μm cell strainer and washed the cells with PBS containing 0.04% BSA. To process HEK293T cells, we collected them at 70–80% confluency, dissociated them with Accutase, and washed them as previously described. We assessed cell concentration and viability using a hemocytometer or an automated cell counter before proceeding. For tumor samples, we performed scRNA-seq on tumors derived from the 1,000-cell and 5,000-cell injection groups. For each 10x lane, we pooled four samples: 1k piece 1, 1k piece 4, 5k piece 1, and 5k piece 4. We encapsulated and barcoded these cells using the 10x Genomics Chromium Single Cell 3′ platform (v3.1 chemistry). During library preparation, we combined all cDNA products from each lane separately (lane 1 and lane 2), and prepared libraries from the pooled cDNAs of each lane. To increase yield, we divided each pooled cDNA sample into three PCR tubes for amplification. We performed library preparation and cDNA amplification according to the manufacturer’s protocol. We then sequenced the libraries on an Illumina platform.

#### Palindromic Feature Weight and Outcome Prediction

To predict feature weights in the 208-target library, we implemented a machine learning pipeline to predict CRISPR gene-editing outcomes using XGBoost with SMOTE to handle class imbalance (Chen and Guestrin 2016; Chawla et al. 2011). We use features including GC content, free energy, and one-hot-encoded positional nucleotides from guide RNA sequences of lengths 17, 18, and 20. For both multi-class edit-type prediction (left/right/both) and binary guide-efficiency identification, the pipeline employs SMOTE to balance minority classes in the training data only, followed by XGBoost classification. Models are evaluated using correlation coefficients, RMSE, precision, recall, F1 scores, and confusion matrices across all guide RNA lengths. Python scripts are available on the GitHub code repository associated with this paper.

#### Lineage Simulation

Lineage simulations were generated with https://github.com/mckennalab/counterfeiter. Cells were divided in a bifurcating pattern for 8 cell divisions, varying the editing balance (left vs right proportion), editing rate per cell generation (from 0.1% to 75%), and the number of integrated targets per cell (5 to 150). Quartet divergence was calculated in R from the ground-truth and reconstructed trees using IQTree2 (Sand et al. 2014; Minh et al. 2020).

#### General Computational Analysis

Code supporting analysis in the paper is available on GitHub at this link: https://github.com/mckennalab/palindrome_base_editing. In brief, lineage barcodes were processed with Clique (https://github.com/mckennalab/clique) using YAML configuration files available in the palindrome GitHub. Further analysis was performed using custom R and Python scripts that invoked other computational tools. These scripts are available on GitHub, including code to reproduce the main figures in the paper.

## Acknowledgements

We thank the members of the McKenna lab for discussion and support. This work was supported by the Dartmouth Cancer Center and the shared resources supported by NCI Cancer Center Support Grant 5P30CA023108. We especially thank Fred W. Kolling IV, Laurent Perreard, Carol Ringelberg, and Elizabeth Sergison of the GMBSR for their guidance and expertise in sequencing. FACS was carried out by Gary Ward in DartLab, the Immune Monitoring and Flow Cytometry Shared Resource at the Dartmouth Cancer Center (RRID: SCR_019165). We also acknowledge the Dartmouth Cancer Center Irradiation, Imaging, Microscopy, and Animal Cancer Models Shared Resource (RRID:SCR_023478), and specifically Jennifer Fields for her help with animal experiment strategy. All Illumina sequencing was carried out in the Genomics and Molecular Biology Shared Resource (RRID:SCR_021293), which is additionally supported by NIH S10 (1S10OD030242) awards. Single-cell studies were conducted through the Dartmouth Center for Quantitative Biology in collaboration with the GMBSR with support from NIGMS COBRE (P20GM130454) and NIH S10 (S10OD025235) awards. This work was supported by DP2GM149750, and A.M. is supported by the Pew Biomedical Scholars program and the V Foundation.

## Data Availability

Lineage-recorder sequencing and 10X Chromium data are available in NCBI GEO under accession number GSE327634.

## Supplementary figures and legends

**Supplemental Figure 1:**
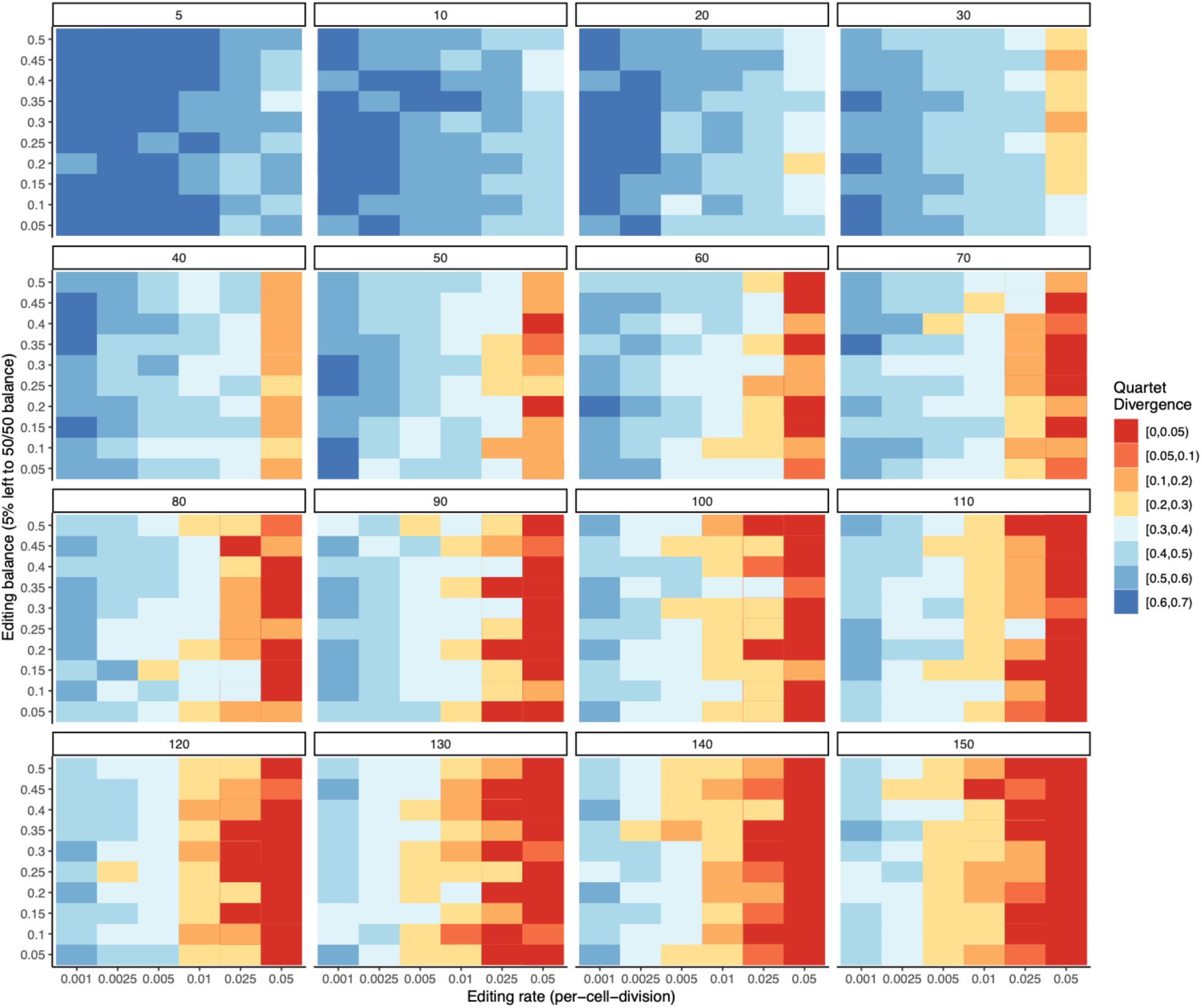
Simulation of PALINCODE editing outcomes. Simulations were run for eight generations, varying the number of target integrations (header box number in each sub-plot), the editing balance (y axis), and the editing rate (x axis). Quartet divergence was then calculated by comparing the known, ground-truth tree with the tree reconstructed from simulated PALINCODE outcomes using IQTree2. Quartet divergence decreases to <5% in as few as 20 integrated targets, with the highest-accuracy trees captured at 5%-25% editing per generation.

**Supplementary Figure 2:**
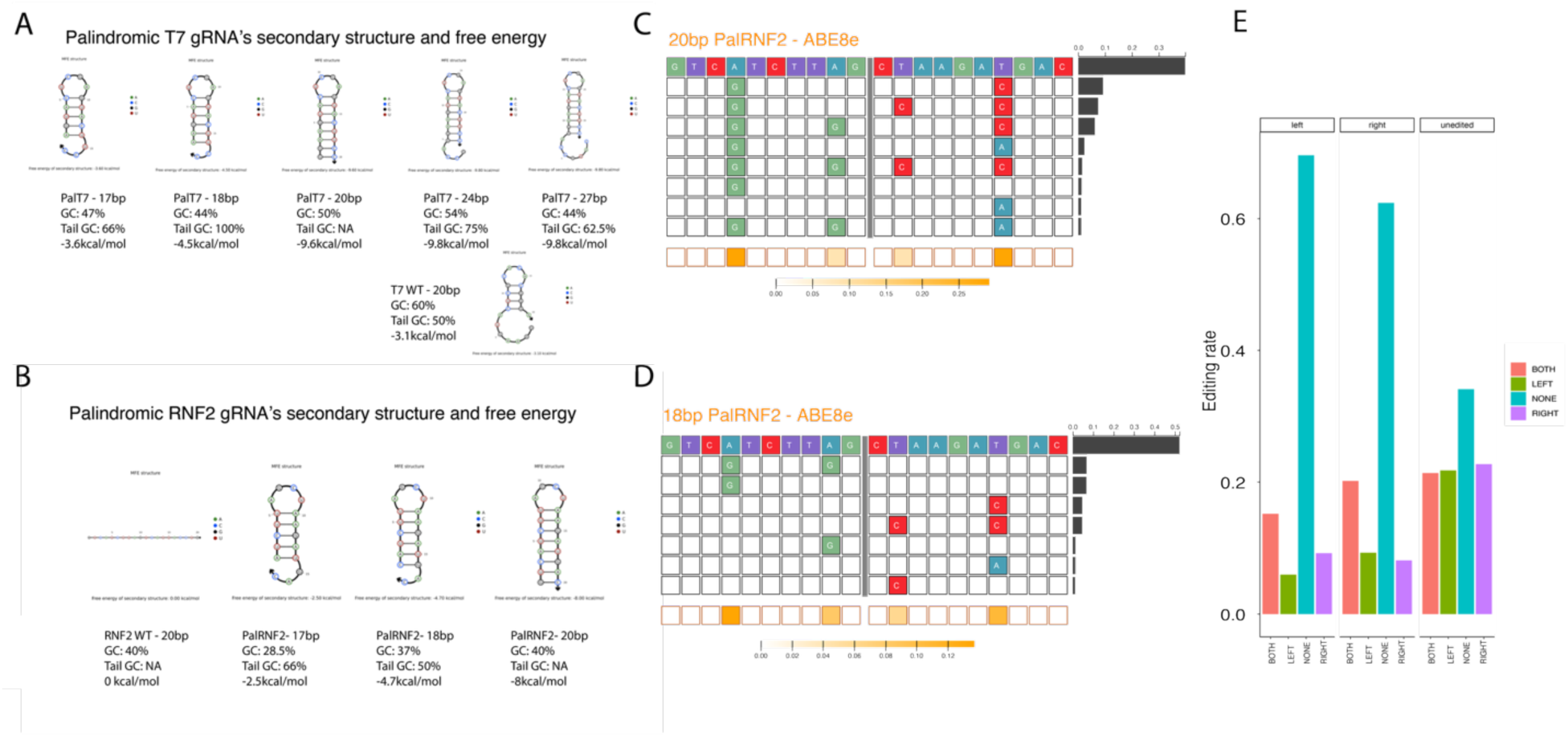
Folding energy effects on editing for both targets PalT7 and PalRNF2. Folding energy and GC content for both the wild-type and the palindromic T7 **(A)** and RNF2 RNA guide sequences **(B)** over a range of sequence lengths. **(C)** Editing outcomes for target PalRNF2 using ABE8e ABE editor with a 20bp guide sequence and **(D)** 18-basepair guide sequence. **(E)** Pre-edited target outcomes for the unedited PalRNF2, left pre-edited, and right pre-edited target libraries with an 18-basepair guide.

**Supplementary Figure 3:**
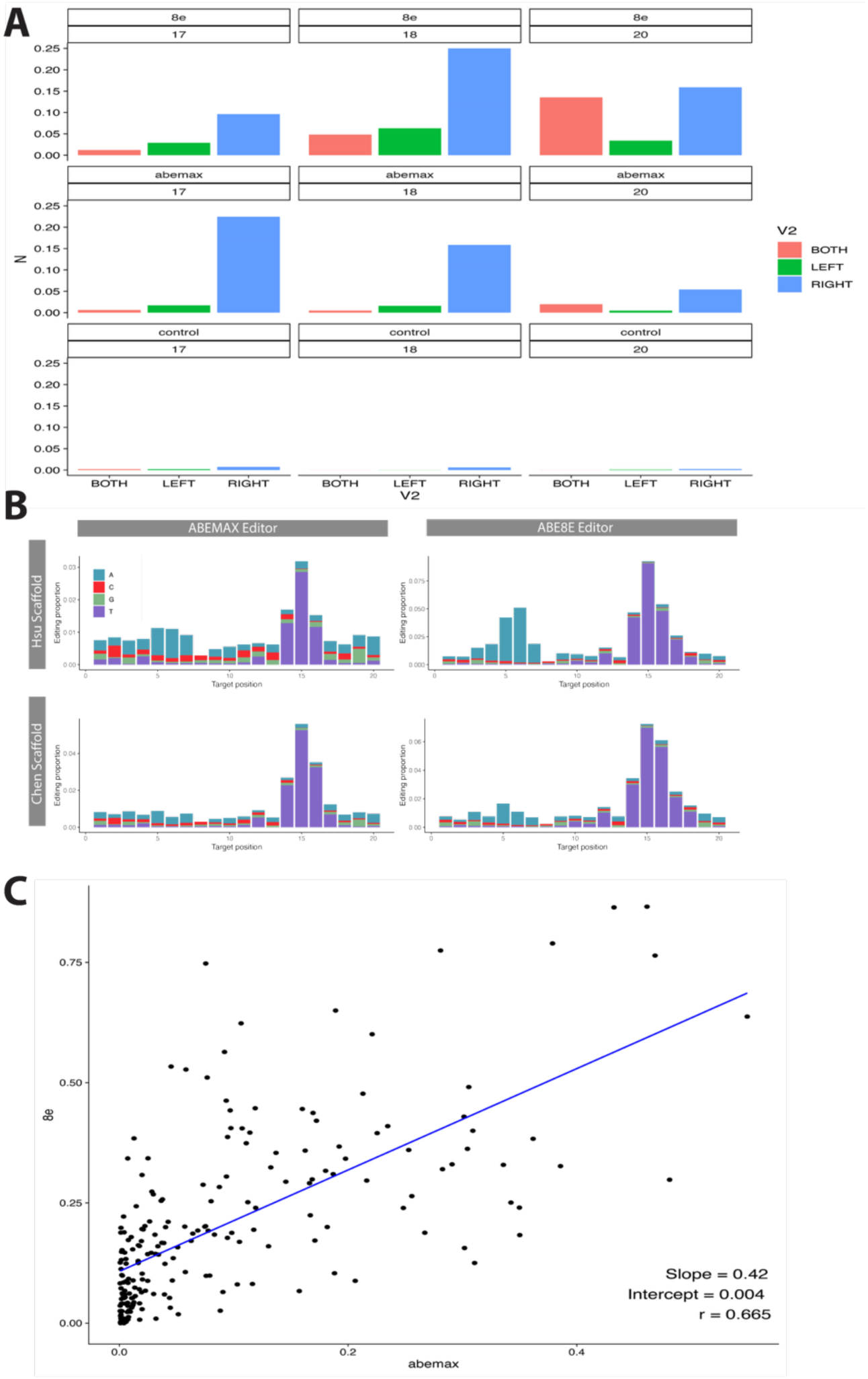
Editing outcomes in oligo library. **(A)** Left, right, and both editing outcomes for different base-editors (ABE 8e and ABEMAX, and control) and three different guide sequence lengths (17, 18, and 20). **(B)** For length 18 guide sequences, comparison of the Hsu and Chan scaffold sequences using both ABE 8e and ABEMAX base editing constructs. **(C)** Correlation between editing rates across all 208 targets for the ABE 8E and ABEMAX editors.

**Supplementary Figure 4:**
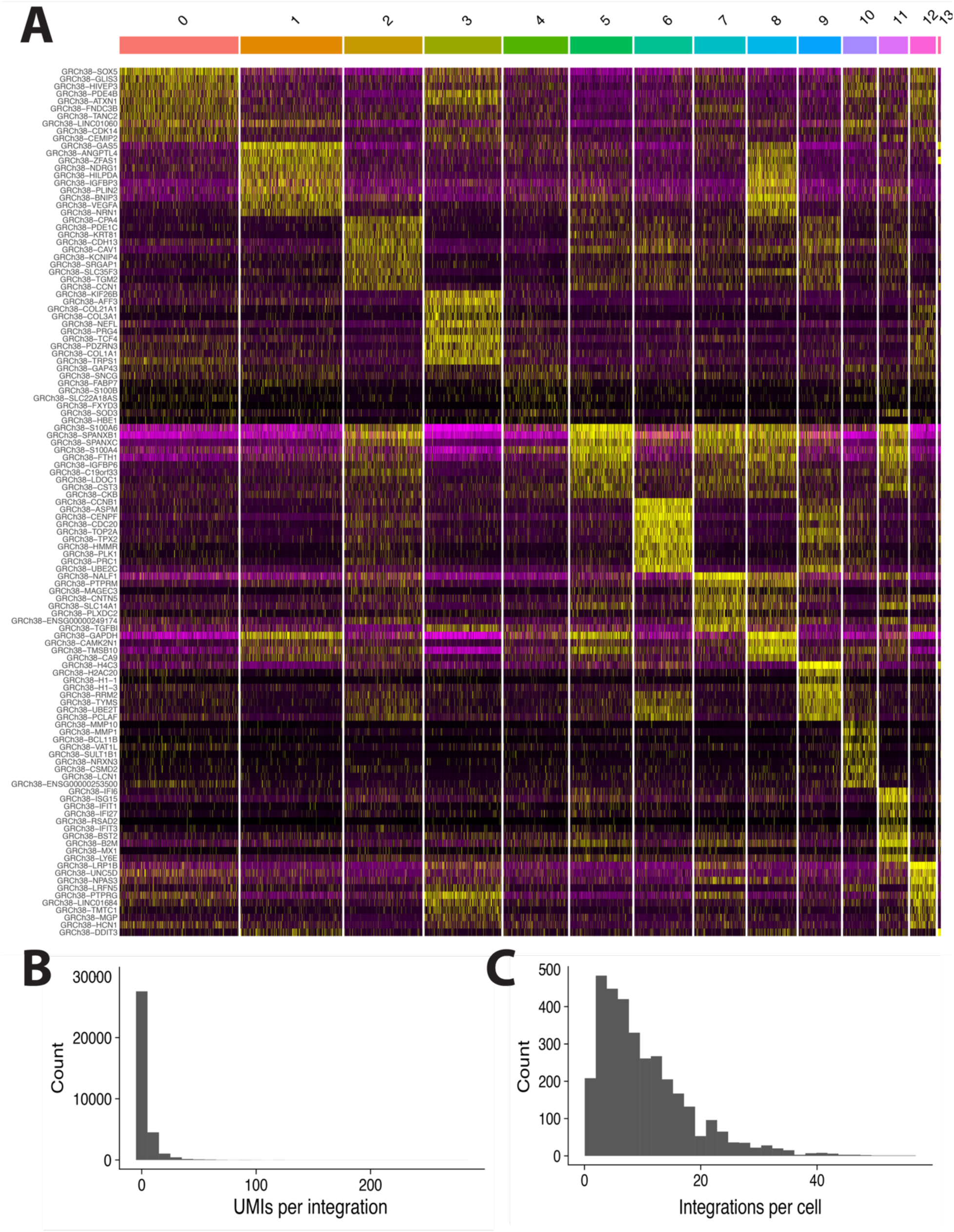
A375 transcriptional classification and lineage capture. **(A)** The top ten significantly differentially expressed genes per single-cell transcriptional cluster are shown in the UMAP in Figure 5B. **(B)** Number of collapsed UMIs per genome integration in the single-cell A375 data set. **(C)** Total number of integrations detected per captured cells.

